# A dual-localized geraniol synthase and a previously unreported cytosolic geranyl pyrophosphatase contribute to geraniol formation in lemongrass

**DOI:** 10.1101/2025.08.29.673030

**Authors:** Priyanka Gupta, Anuj Sharma, Sruthi Mohan, Yash Misra, Ramanathan Natesh, Dinesh A. Nagegowda

**Affiliations:** Molecular Plant Biology and Biotechnology Lab, CSIR-Central Institute of Medicinal and Aromatic Plants Research Centre, Bengaluru - 560065, India; Academy of Scientific and Innovative Research (AcSIR), Ghaziabad – 201002, India; School of Biology and Indian Institute of Science Education and Research Thiruvananthapuram, Maruthamala P.O., Vithura, Thiruvananthapuram – 695551; Center for High-Performance Computing (CHPC), Indian Institute of Science Education and Research Thiruvananthapuram, Maruthamala P.O., Vithura, Thiruvananthapuram – 695551

## Abstract

Geraniol, a major constituent of many essential oils and precursor for geraniol-derived aldehydes and acetates, is produced in plants through the action of a terpene synthase encoding geraniol synthase or via terpene synthase–independent non-canonical pathway involving a Nudix hydrolase and a yet to be discovered pyrophosphatase. Here, we identify a geraniol synthase (*Cymbopogon flexuosus* geraniol synthase; CfGES) that catalyzes the conversion of geranyl pyrophosphate (GPP) (also known as geranly diphosphate) to geraniol, as well as a previously unreported geranyl pyrophosphatase (CfG(P)Pase) that acts on both GPP and geranyl monophosphate (GP), albeit with different efficiencies, to generate geraniol. Virus-induced gene silencing of *CfGES* or *CfG(P)Pase* resulted in a substantial reduction in geraniol and its immediate product citral in lemongrass leaves. Conversely, transient overexpression of *CfGES* or *CfG(P)Pase* resulted in enhanced production of geraniol and citral in lemon balm (*Melissa officinalis*) leaves, as well as geraniol, citronellol, and citral in rose (*Rosa damascena*) flower petals. Subcellular localization studies revealed that while CfGES exhibited dual cytosolic/plastidial localization, CfG(P)Pase was localized to the cytosol. This localization pattern was further supported by the significantly higher geraniol-forming activity observed in the purified cytosolic protein fraction compared to the chloroplast fraction. Our study uncovers the missing step in cytosolic geraniol formation via a TPS-independent non-canonical route and demonstrates that geraniol, required for citral production in lemongrass, is synthesized by cytosolic CfG(P)Pase and cytosol-/plastid-localized CfGES.

## Introduction

Aromatic grasses of the *Cymbopogon* genus, unique members of the Poaceae family, produce substantial amounts of monoterpene-rich essential oils. The major constituents of these essential oils consist of the monoterpene alcohol geraniol and/or its aldehyde or ester derivatives. For example, palmarosa (*Cymbopogon martinii*) produces geraniol-rich essential oil, lemongrass (*Cymbopogon flexuosus*) predominantly accumulates essential oil composed of geraniol aldehydes (citral), and citronella grass (*Cymbopogon winterianus*) accumulates a mixed-type oil consisting of alcohol and aldehyde constituents. Lemongrass primarily produces citral, which constitutes 70–80% of its essential oil. Citral, composed of neral and geranial isomers, imparts a distinctive lemon-like fragrance to the oil. Due to its aromatic properties, lemongrass oil is widely used in perfumery, cosmetics, and as a flavoring agent in soft drinks. Additionally, it serves as a key precursor in the synthesis of vitamin A through ionones (Majewska et al., 2019). Beyond industrial uses of its essential oil, lemongrass is widely used in herbal teas, other non-alcoholic beverages, and baked foods. While geraniol is a commonly found monoterpene alcohol in volatile profile of many plants, in lemongrass it serves as precursor for citral (Gupta et al., 2024). However, the enzyme(s) involved in geraniol formation have not been characterized in aromatic grasses including lemongrass. We recently demonstrated that citral formation in lemongrass is catalyzed by two phylogenetically distant enzymes: a dual-localized CfADH1 (present in both the cytosol and plastids) and a cytosol-localized CfAKR2b. Both CfADH1 and CfAKR2b utilize geraniol as a substrate to produce citral. Additional findings from our previous study, including subcellular fractionation, pathway-specific inhibitor feeding, and *in planta* studies, indicated that geraniol needed for citral formation is derived from substrates synthesized via both cytosolic and plastidial pathways (Gupta et al., 2024). If this scenario holds true, it necessitates the presence of geraniol-forming enzymes in both the cytosol and plastids in lemongrass.

It has been shown that geraniol synthase (GES), a member of the terpene synthase family, catalyzes the formation of geraniol using GPP as a substrate mostly in plastids or in some cases in the cytosol (Iijima et al., 2004). Additionally, emerging evidence shows the existence of a non-canonical TPS-independent route for geraniol biosynthesis, mediated by a novel class of enzyme in certain plant species. It was first reported in rose that the scent component of petals, geraniol, is derived via a Nudix (nucleoside diphosphate-linked moiety X hydrolases) enzyme, which converts GPP to GP. Subsequently, GP was proposed to be converted to its alcoholic form geraniol by a yet to be discovered pyrophosphatase. After this discovery in rose, few more studies followed and reported similar Nudix enzymes involved in geraniol and geranyl β-primeveroside biosynthesis in geranium and tea, respectively (Bergman et al., 2021; Zhou et al., 2022). Nudix hydrolase was also reported to be involved in borneol biosynthesis by hydrolysis of bornyl diphosphate (BPP) to generate bornyl phosphate (BP) in *Wurfbainia villosa* (Yang et al., 2024). Therefore, we hypothesized that geraniol required for citral biosynthesis in lemongrass could be produced through the action of GES, NUDX/pyrophosphatase, or both (Figure S1).

## Results and Discussion

Based on transcriptome and gene expression analyses, we had previously identified a putative terpene synthase (*CfTPS1*) and a putative PPase (*CfPPase1*) in lemongrass (Meena et al., 2016). Before taking up functional characterization of these genes, we first wanted to validate the geraniol forming activity in leaf crude protein extracts of lemongrass. For this, crude protein extracts were subjected to enzyme assay by incubating with GPP and GP in presence of cofactor MgCl_2_ and MnCl_2_. Analysis of assay products by GC/GC-MS showed that both substrates yielded a significant peak for the geraniol product, implying that the crude extract possesses enzymatic activity capable of converting both GPP and GP into geraniol (Figure 1a). Further quantification of geraniol-forming ability in leaf extracts revealed comparable activity levels when GPP or GP was used as a substrate (Figure 1b). This indicates that geraniol biosynthesis in lemongrass occurs through both TPS-dependent and TPS-independent pathways. Next, to determine whether phosphatase activity is involved in the enzymatic conversion of GPP or GP, we incubated the crude protein extracts with GPP or GP along with a phosphatase inhibitor (PI) (PhosStop, Sigma Aldrich). GC analysis of the assay products revealed a significant reduction in geraniol formation in the PI-treated samples compared to the control (Figure 1 c & e). Notably, geraniol-forming activity exhibited greater inhibition when GPP (79.23%) was used as a substrate compared to GP (63%) (Figure 1 d & f). This substantiates that the crude extract contains either a GES, which converts GPP into geraniol through diphosphohydrolase activity, or a pyrophosphatase, which converts GPP and/or GP into geraniol via phosphatase activity. These results together with our previous identification of *CfTPS1* and *CfPPase1* reinforced us to determine the potential role of these genes in geraniol formation. Hence, we next investigated the expression of these candidate genes across various developmental phases (1 month, 2 months, 3 months, and harvesting) of lemongrass leaves, which revealed that the expression of *CfTPS1* exhibits a gradual increase from the 1-month stage to the harvesting phase, implying its sustained participation in geraniol biosynthesis as the plant undergoes maturation (Figure 1g). In contrast, *CfPPase1* transcript levels peaked at the 2-month stage and subsequently declined (Figure 1h).

**Figure 1.**
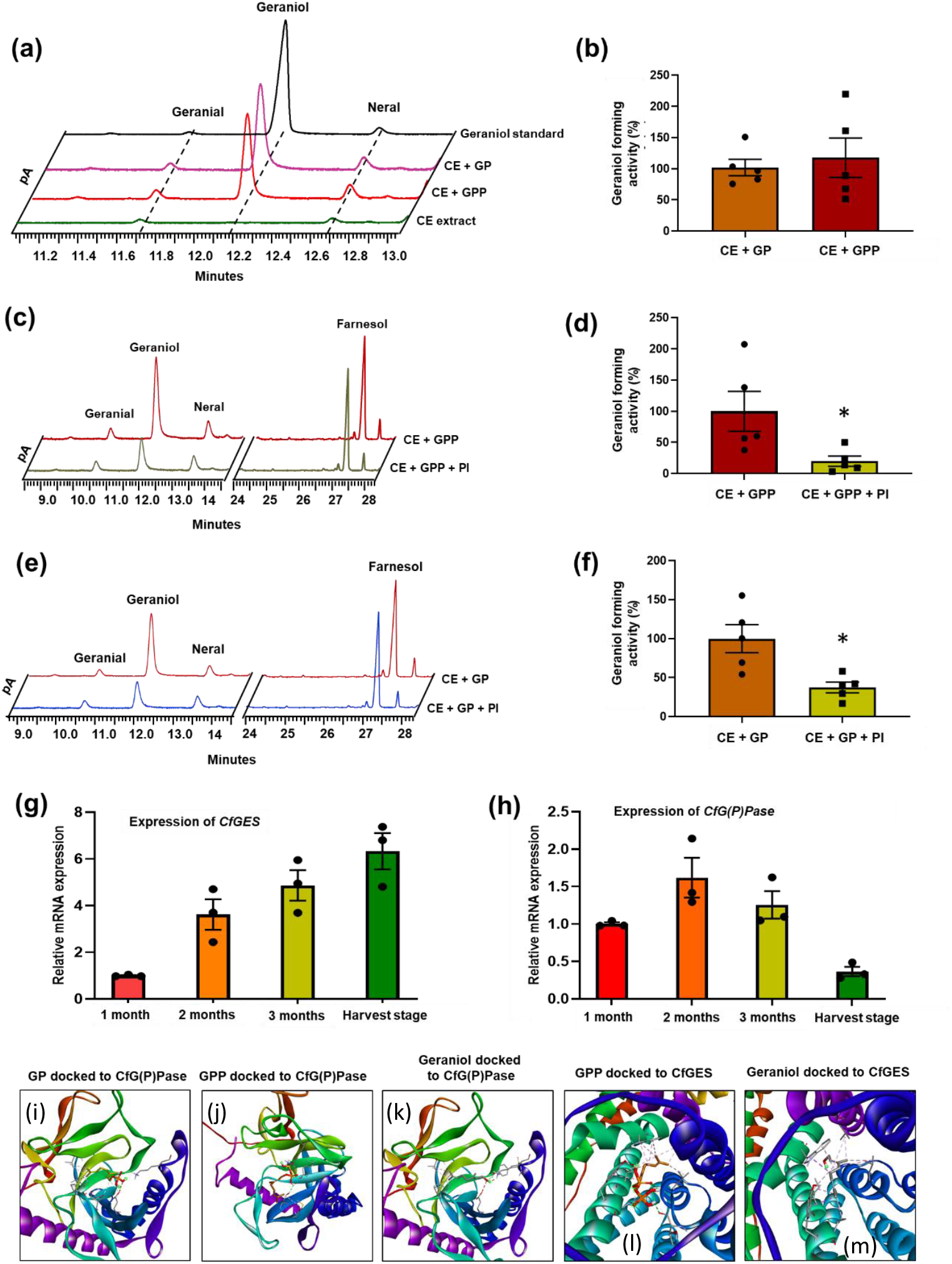
Geraniol-forming activity in lemongrass, along with gene expression analysis and molecular docking of candidate genes and their encoded proteins. **a**, Determination of presence of geraniol-forming activity in crude total protein extracts of lemongrass leaves. The activity assay was carried out by incubating the leaf crude protein extract in the presence or absence (control) of GPP or GP as the substrate and NADP cofactor, with a hexane overlay. Reaction products extracted into the hexane fraction were analysed by GC/GC-MS and the products formed in the assay were compared to the authentic geraniol and citral standards. Crude extract without any substrates served as a control. **b**, Quantification of % conversion of GPP and GP into geraniol. Peak area of geraniol formed in the assay was quantified in relation to the IS. **c&e**, Analysis of inhibitory effect of phosphatase inhibitor on geraniol-forming ability of lemongrass leaf crude protein. About 25-50 ug of total leaf extract was incubated with GPP or GP with or without alkaline phosphatase inhibitor (50 µM). **d&f**, Quantification of % reduction in geraniol-forming ability of lemongrass leaf protein extract in the presence of PI. **g&h**, Reverse transcriptase-quantitative polymerase chain reaction (RT-qPCR) analysis showing the relative expression of *CfGES* and *CfG(P)Pase* in different developmental stages of lemongrass. Total RNA isolated from leaves was used for RT-qPCR analysis, and the expression level of genes was normalized to *CfEF1α* endogenous control and was set to 1. **i to m**, Molecular docking of CfG(P)Pase with GP ((i), GPP (j), and geraniol (k), and CfGES with GPP (l) and geraniol (m).

This expression pattern implies a more significant role for *CfPPase1* during earlier developmental phases, with diminishing transcriptional activity as development proceeds towards maturity. Collectively, these differential expression profiles highlight the dynamic temporal regulation of key enzymatic components in the geraniol biosynthetic pathway during lemongrass leaf development.

Prior to taking up functional characterization, we adopted computational methods to investigate the binding affinities and interactions between CfG(P)Pase with GP, GPP, geraniol and FPP, and CfGES (referred to as CfTPS1) with GPP and geraniol (Figure 1i-m & Figure S2 a-h). The results indicated that CfG(P)Pase interacts with GPP via four hydrogen bonds (Figure S2 e), while interactions with GP and geraniol are primarily mediated by Van der Waals forces with only one hydrogen bond formed with GP (Figure S2 d) and geraniol (Figure S2 c). Although FPP showed a similar binding energy to GPP, this result is likely due to the presence of the pyrophosphate moiety in the molecule (Figure S2 f). Docking result showed that CfGES has a binding affinity of −7.9 kcal/mol with GPP, forming one hydrogen bond with Threonine 415. Also, it was noted that CfGES has better binding affinities for GPP compared to Cf(G)PPase (Figure S2). The indication of GES- and PPase-dependent geraniol formation in the crude extract, the correlation between the expression of *CfGES* and *Cf(G)PPase* with geraniol accumulation, and the docking analysis showing favourable binding energies of both enzymes with GPP and GP collectively prompted us to undertake biochemical characterization to determine their enzymatic activity. To assess the biochemical nature of CfTPS1 and CfPPase1, we first cloned them into an *E. coli* expression vector, generated recombinant proteins, and purified them using affinity chromatography. Biochemical assay with GPP as the substrate, MgCl_2_ and MnCl_2_ as cofactor, followed by GC-MS analysis, revealed that the purified recombinant CfTPS1 catalyzed the formation of geraniol as the sole product. In contrast, CfTPS1 did not produce any detectable product when FPP was used as the substrate, indicating its specificity for monoterpene synthase activity, and hence CfTPS1 is hereafter referred as CfGES (Figure 2 a & c). The geraniol synthase activity of CfGES, characterized by its specificity for GPP and exclusive production of geraniol, closely aligns with the functional profiles of previously characterized geraniol synthases from other plant species. These include CrGES from *Catharanthus roseus* (Madagascar periwinkle), DoGES from *Dendrobium officinale*, ObGES from *Ocimum basilicum*, RdGES from *Rosa damascena*, and CtGES from *Cinnamomum tenuipilum* (Simkin et al. 2013; Dong et al. 2013; Chen et al. 2016; Zhao et al. 2020; Zhou et al. 2022; Jiang et al. 2023). The similarity in substrate specificity and product formation reinforces the function of CfGES as a *bona fide* geraniol synthase. Similar to assay with CfGES, the assay using recombinant CfPPase and GPP as substrate showed formation of geraniol as the sole product. In addition, CfPPase was able to catalyze conversion of GP to geraniol in a much efficient manner than that of GPP to geraniol. The assay using purified protein from empty vector-transformed *E. coli* did not show geraniol formation with either GPP or GP as a substrate, ruling out the involvement of endogenous *E. coli* phosphatases. Furthermore, CfPPase1 could not hydrolyze FPP to farnesol confirming its specific GP/GPP-hydrolyzing and geraniol-forming activity and hence CfPPase1 is hereafter referred as CfG(P)Pase (Figure 2 b&d). To further ascertain the pyrophosphatase activity, an enzyme assay was conducted in the presence of a phosphatase inhibitor (PI), using GPP and GP as substrates. The geraniol-forming activity of CfG(P)Pase was inhibited by the PI with both substrates. Notably, the inhibition was more pronounced with GPP (90%) compared to GP (69%), aligning with the inhibition patterns of pyrophosphatase activity observed in crude leaf extracts treated with PI (Figure S3). The stronger inhibition observed with GPP is conceivable as CfG(P)Pase can convert both GPP and GP into geraniol. So far, no gene encoding a PPase involved in geraniol formation has been identified in plants. However, PPase activity in leaf crude extract of rice has been reported to be involved in the formation of farnesol and geranylgeraniol (Nah et al., 2001).

**Figure 2.**
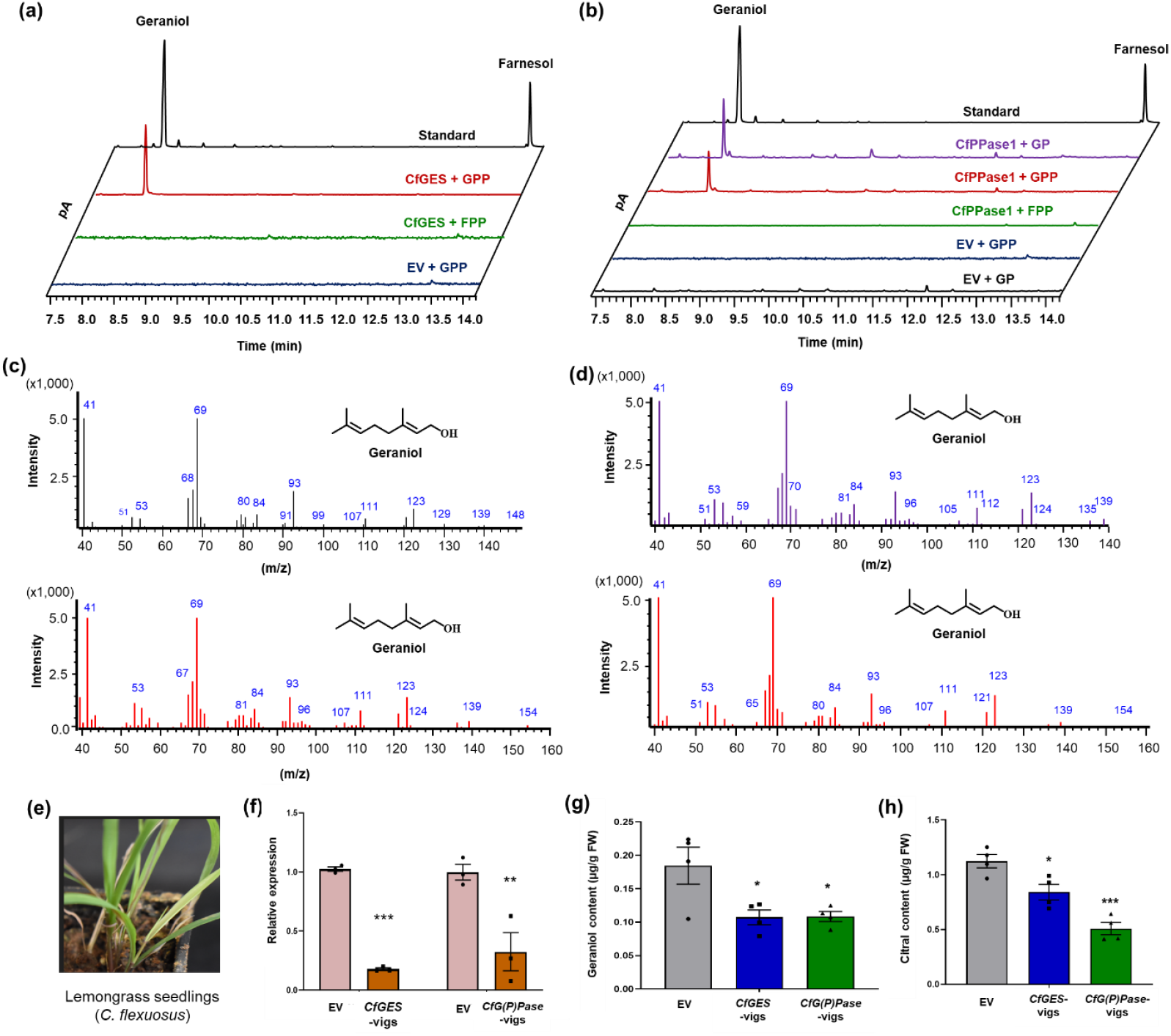
Biochemical and *in planta* proof for the involvement of CfGES and CfG(P)Pase in geraniol biosynthesis. **a**, Determination of reaction products formed by recombinant CfGES. Gas Chromatography-Mass Spectrometry (GC-MS) profile showing the products formed by CfGES from GPP. The enzyme activity assay was carried out using the purified recombinant CfGES in the presence of GPP or FPP as a substrate and MgCl_2_ cofactor, with a hexane overlay. Assay with purified protein from *E. coli* transformed with empty vector (EV) in presence of GPP and MgCl_2_ served as control. b, Analysis of reaction products formed by recombinant CfG(P)Pase. GC-MS profile showing the products formed by CfG(P)Pase from GP and GPP. The enzyme activity assay was carried out using the purified recombinant CfG(P)Pase in the presence of GP or GPP or FPP as a substrate and MgCl_2_ cofactor, with a hexane overlay. Assay with purified protein from *E. coli* transformed with EV in presence of GP or GPP and MgCl_2_ served as control. In both a & b, reaction products formed in the assay and unreacted substrate were extracted into the hexane fraction and analysed by GC-MS. The products formed in the assay was confirmed by comparing the retention time of authentic citral standard. **c & d**, Mass spectra of product peaks of the assay. The products formed in the assay were further confirmed by comparing the mass spectra of authentic geraniol standard. **e**, Phenotype of leaves infected with pTRV2::*CfPDS* 30 days post agro-infection. **f**, Reverse transcriptase-quantitative polymerase chain reaction (RT-qPCR) analysis showing the relative expression of *CfGES*, and *CfG(P)Pase* in respective VIGS samples of lemongrass. Total RNA isolated from EV control, *CfGES*-vigs, and *CfG(P)Pase*-vigs leaves was used for RT-qPCR analysis. The expression level of genes was normalized to *CfEF1α* endogenous control and was set to 1 in EV control to determine the relative reduction of transcripts in *CfGES*-vigs and *CfG(P)Pase*-vigs leaves. **g&h**, Relative level of geraniol (g) and citral (h) in *CfGES*-vigs and *CfG(P)Pase*-vigs leaves in comparison to EV control. In all experiments, volatiles were extracted using hexane containing internal standard (IS) *E,E*-farnesol, and subjected to gas chromatography (GC) analysis. The amount of geraniol and citral was quantified in relation to the IS. The results shown are from four independent experiments indicated by data points. Statistical analysis was performed by two-way ANOVA (RT-qPCR) and one way ANOVA (metabolites) using GraphPad Prism 8: ***, *P*<0.001. Error bars indicate mean ± SE.

Since both CfGES and CfG(P)Pase could form geraniol in *in vitro* assays, we next wanted to establish definitively their *in planta* role in geraniol biosynthesis. First, we downregulated the expression of *CfGES* and *CfG(P)Pase* in lemongrass leaves by virus-induced gene silencing (VIGS) (Figure 2e), which was established for the first time in our recent study (Gupta et al., 2024). Specific silencing of *CfGES* led to 82% decreased transcripts with a corresponding decrease in the levels of geraniol and citral by 45% and 25%, respectively (Figure 2 f – h & S4a). This indicated that *CfGES* forms geraniol *in planta* thereby providing substrate for citral biosynthesis. Similarly, VIGS of *CfG(P)Pase* showed a 68% reduction in its transcript levels resulting in a corresponding reduction in geraniol and citral levels by 46% and 55%, respectively (Figure 2 f -h & S4a).

Gene expression analysis followed by metabolite quantification of leaves subjected to VIGS clearly showed the role of both *CfGES* and *CfG(P)Pase* in geraniol formation in lemongrass. To further verify this finding, we adopted transient overexpression approach. Since lemongrass leaves are not amenable for agroinfiltration, lemon balm (*Melissa officinalis*), which also produces similar terpene profile to that of lemongrass, was used for transient overexpression of *CfGES* and *CfG(P)Pase* using vacuum infiltration. Indeed, this method was successfully utilized in our recent study to show the role of CfADH1 and CfAKR2b in citral formation (Gupta et al., 2024). Transient overexpression of *CfGES* and *CfG(P)Pase* in lemon balm leaves was confirmed by semi-quantitative RT-PCR using gene specific primers (Figure 3 a & b). Subsequent metabolite analysis showed 1.26-fold and 1.58-fold increase of geraniol and citral in *CfGES* overexpressing leaves in comparison to empty vector control. In a similar manner, overexpression of *CfG(P)Pase* resulted in 1.43-fold and 1.46-fold increase in geraniol and citral content, respectively (Figure 3 c, d & S4b). To validate the genes in another heterologous platform, we utilized Rose (*Rosa damascena*) petals as they are a source of geraniol and minor amounts of citral and citronellol. It has been shown that geraniol is formed in rose petals via non-TPS route by a Nudix hydrolase (Figure S5). Hence, to determine if overexpression of *CfGES* and *CfG(P)Pase* has any effect on monoterpenes, we performed transient overexpression of *CfGES* and *CfG(P)Pase* by agroinfiltration of Rose petals. Analysis of gene expression by semi-quantitative RT-PCR after 48 hours showed the expression of corresponding genes (Figure 3 e & f). Subsequent volatile analysis using headspace GC-MS and quantification showed an increase in geraniol in both *CfGES* and *CfG(P)Ppase* infiltrated petals compared to empty vector control (Figure 3h). The increase in geraniol accumulation was followed by enhanced levels of citral and citronellol (Figure 3g & S5c), indicating that the excess geraniol produced due to the overexpression of *CfGES* and *CfG(P)Pase* in rose petals was utilized by endogenous enzymes for its conversion into citral and citronellol. The enhanced production of geraniol and citral in lemon balm, along with geraniol, citral, and citronellol in rose petals, in the *CfGES* and *CfG(P)Pase* overexpression background not only provided additional proof of their role in geraniol formation but also provided strong evidence for the presence of a GPP pool in both plastids and the cytosol. Moreover, it highlights the coexistence of both canonical TPS-dependent and non-canonical TPS-independent pathways for geraniol biosynthesis in lemon balm and rose similar to lemongrass. Indeed, our search for candidate genes indicated the presence of putative GES and PPase in lemon balm with a similarity of 41% and 73% and Rose (*Rosa chinensis*) with a similarity of 32% and 83% respectively, (Figure S6 & S7). The strong evidence from both VIGS in lemongrass and overexpression studies in lemon balm and rose clearly establishes the *in planta* role of *CfGES* and *CfG(P)Ppase* in geraniol formation. This further suggests that these two enzymes function upstream of CfADH1 and CfAKR2b, which are responsible for catalyzing the conversion of geraniol formed by CfGES and GfG(P)Pase into citral.

**Figure 3.**
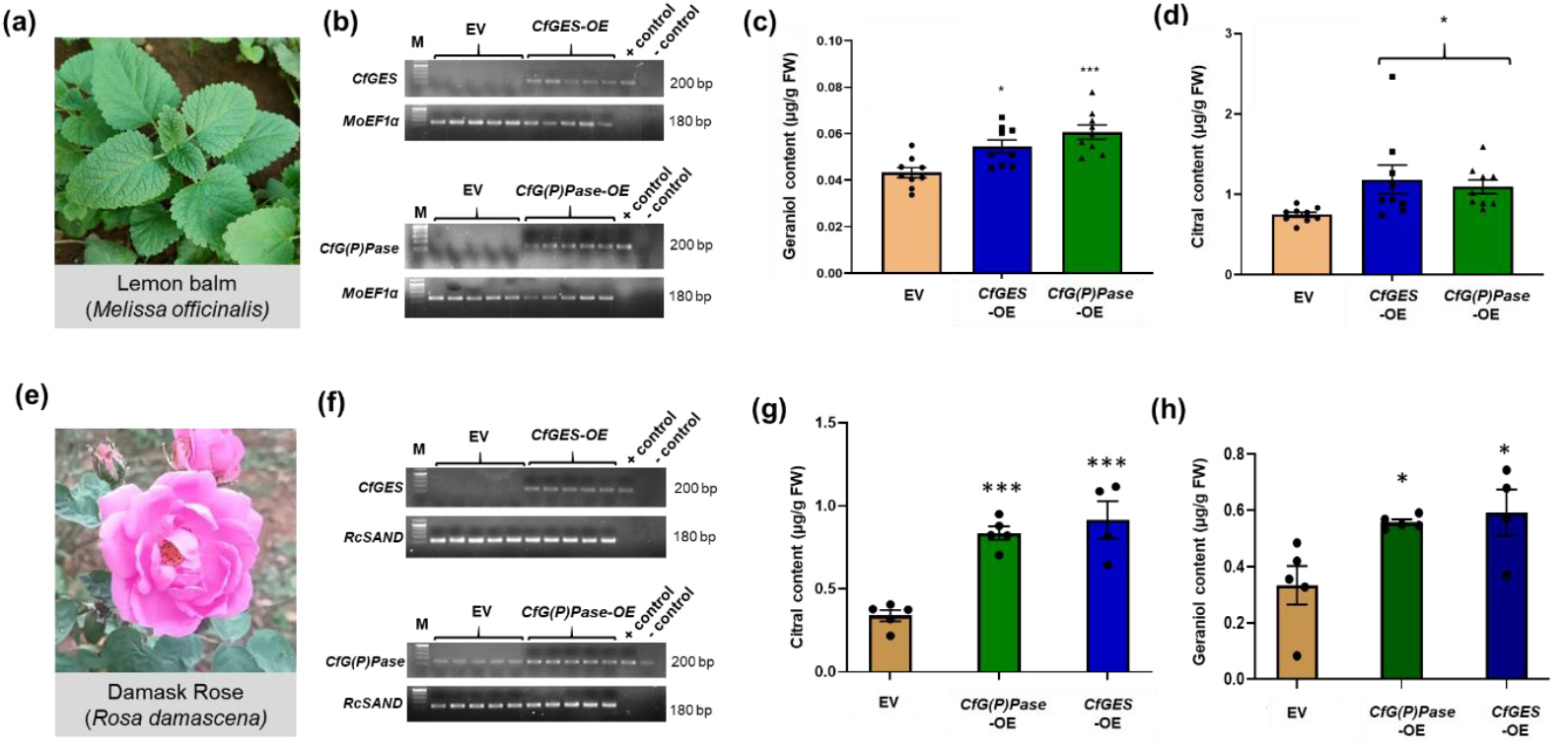
Transient expression of *CfGES* and *CfG(P)Pase* and its effect on geraniol biosynthesis and citral content in lemon balm (*Melissa officinalis*) and Damask Rose (*Rosa damascena*). (a & e), Representative image of the lemon balm plant and damask rose flower used for transient expression. (b & f) Semi-quantitative reverse transcriptase polymerase chain reaction analysis showing the band intensity of *CfGES* and *CfG(P)Pase* in respective overexpression samples. Total RNA isolated from empty vector (EV control), *CfGES* −OE, and *CfG(P)Pase* −OE leaves was used for semi-quantitative RT-PCR analysis. Volatiles were extracted using hexane containing internal standard (IS) E,E-farnesol and subjected to gas chromatography (GC) analysis. (c & d) Determination of geraniol and citral content in lemon balm leaf tissues infiltrated with Agrobacterium harboring EV control, *CfGES* −OE, and *CfG(P)Pase* −OE constructs. (g & h) Determination of geraniol and citral content in damask rose tissues infiltrated with Agrobacterium harboring EV control, *CfGES* −OE, and *CfG(P)Pase* −OE constructs. The amount of geraniol and citral was quantified in relation to the IS. The results shown are from 5 (metabolites) independent experiments indicated by data points. Statistical analysis was performed by two-way ANOVA (RT-qPCR) and one-way ANOVA (metabolites) using GraphPad Prism 8: \**P* < 0.05; \*\**P* < 0.002; \*\*\**P* < 0.001. Error bars indicate mean± SE.

Geraniol is a monoterpene and hence is traditionally known to be synthesized from GPP derived via the plastidial MEP pathway. Accordingly, GESs are known to be localized in plastids as has been shown for GESs from *C. roseus*, DoGES, VoGES, OsTPS21, and PlGES (Iijima et al., 2004; Dong et al., 2013; Simkin et al., 2013; Blerot et al., 2018; Zhao et al., 2020). However, GES from *Lippia dulcis* was reported to have both cytosolic and plastidial localization (Dong et al., 2013). More recently, a cytosol-localized GES was identified in *Epiphyllum oxypetalum* (Zhang et al., 2025). Furthermore, in rose and geranium, geraniol has been shown to be synthesized in the cytosol by cytosolic NUDX, with the required GPP in rose being produced by a cytosolic bifunctional geranyl/farnesyl diphosphate synthase (RcG/FPPS1) (Bergman et al., 2021; Conart et al., 2023). In our previous study, CfADH1 was shown to have dual localization in both the cytosol and plastids, while CfAKR2b was exclusively localized in the cytosol, indicating that geraniol, the substrate of these enzymes, is produced in both compartments (Gupta et al., 2024). Hence, we next investigated the subcellular localization of CfGES and CfG(P)Ppase so as to determine if they are functioning upstream of CfADH1 and CfAKR2b. Since *in silico* analysis using different prediction tools did not provide clarity on their subcellular location (Table S1), we used YFP fusion proteins to check their *in planta* subcellular compartmental location by transient overexpression in *N. benthamiana* leaves. Confocal microscopy analysis of fluorescence in CfGES-YFP infiltrated leaves revealed a diffuse signal throughout the cell, similar to the fluorescence observed in the cytosolic YFP control. Additionally, CfGES-YFP infiltrated leaves exhibited distinct punctate signals that overlapped with chlorophyll autofluorescence (AF), providing clear evidence of dual cytosolic and plastidial localization (Figure 4a & S8a). In contrast, CfG(G)Ppase-YFP fluorescence was exclusively associated with the cytoplasm, resembling the fluorescence observed in the YFP control sample. Unlike CfGES-YFP, no signal overlap with chlorophyll autofluorescence was detected, confirming the strictly cytosolic nature of CfG(P)Ppase (Figure 4a & S8a). The dual localization of CfGES was similar to localization pattern of CfADH1, whereas the localization of CfG(P)Pase was akin to localization of CfAKR2b, indicating their respective upstream roles in citral biosynthesis. To further validate the subcellular localization findings from confocal microscopy of fusion-YFP proteins, we performed subcellular fractionation of lemongrass leaf protein extracts. The purity of the isolated protein fractions was confirmed using marker enzyme assays, and chloroplast integrity was verified through confocal imaging (Figure S8 b & c). After checking the purity, proteins from the purified subcellular fractions were then subjected to enzyme assays with GPP as substrates, and the resulting products were analyzed and quantified by GC. The analysis revealed that geraniol-forming activity was distributed between the cytosolic and chloroplast fractions, accounting for 65.78% and 23.67% of the total leaf protein activity, respectively (Figure 4 b & c). This suggests that the higher activity in the cytosolic fraction results from the combined contributions of the cytosolic portion of dual-localized CfGES and cytosolic CfG(P)Pase, while the activity in the chloroplast fraction is solely attributable to the plastidial portion of CfGES.

**Figure 4.**
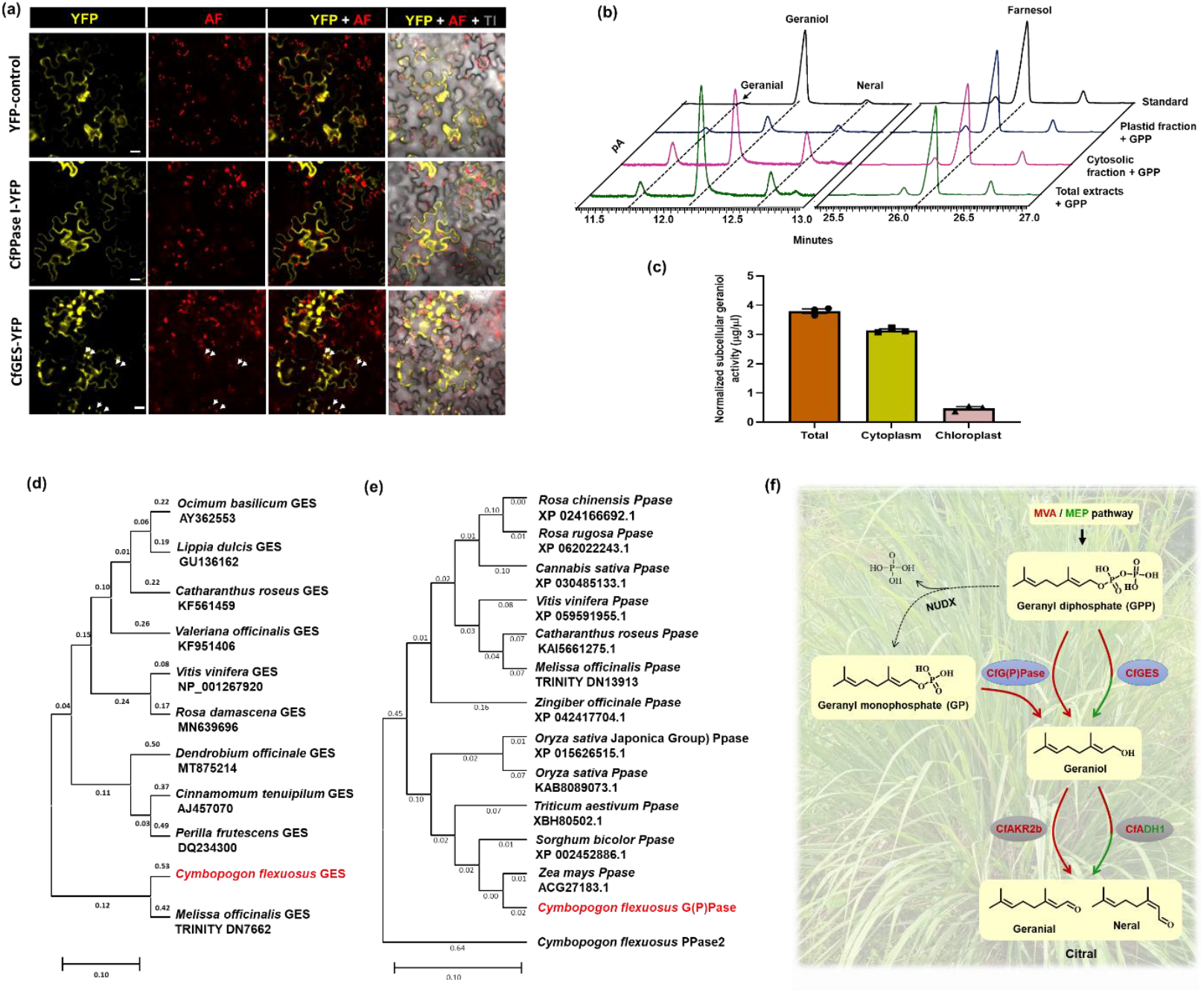
Determination of subcellular localization of CfGES and CfG(P)Pase, and geraniol formation in lemongrass, phylogenetic analysis, and summary of discovered geraniol-pathway. **a**, Confocal laser scanning microscopy of *Nicotiana benthamiana* leaves transformed with CfGES-YFP and CfG(P)Pase-YFP expression constructs. YFP fluorescence and chlorophyll autoflourescence are shown in YFP and AF columns, respectively. The third vertical panel shows merged image of YFP and AF, and the fourth and the rightmost panel shows merged YFP, AF, and transmission image. The arrows indicate chloroplasts. Scale bar: 20 µm. **b & c**, Representative chromatograms and geraniol forming activity in total leaf protein, and cytosolic and plastidic subcellular fractions. In experiments of **b to c**, total crude protein was extracted from lemongrass leaf and was subjected to subcellular fractionation to obtain cytosolic and plastid fractions. Proteins were further desalted and subjected to assay in presence of GPP or GP and Mgcl_2_ and Mncl_2_ as cofactor. Volatiles were extracted using hexane containing internal standard (IS) *E,E*-farnesol, and subjected to gas chromatography (GC) analysis for quantification. The results shown are from three independent experiments indicated by data points. Statistical analysis was performed by simple *t*-test using GraphPad Prism 8.0: \*\*\**P*<0.001. Error bars indicate mean ± SE. **e&f**, Phylogenetic analysis of CfGES (d) and CfG(P)Pase (e) with characterized GESs and putative PPase orthologs of other plants. Phylogenetic tree was constructed using the neighbor-joining method with bootstrap value of 1,000 runs using MEGA11 software (Tamura et al., 2021). Branch lengths denote the genetic distance (number of substitutions per site), with the scale bar indicating 0.20 substitutions per site. **f**, Overview of geraniol biosynthesis in lemongrass. While the dual cytosol/plastid localized CfGES forms geraniol by utilizing GPP made in plastid or cytosol, the cytosolic CfG(P)Pase forms geraniol by utilizing GP or GPP or both in cytosol. Abbreviations: GPP, geranyl diphosphate; Geranyl monophosphate (GP); MEP, methylerythritol phosphate; MVA, mevalonic acid.

So far, GESs have been functionally characterized primarily in dicot plant species, with only two instances reported from monocots (Zhao et al., 2020). In contrast, no PPases involved in geraniol biosynthesis had been identified prior to our study. To explore the evolutionary significance and functional relationships, we conducted sequence analysis and phylogenetic analysis of CfGES with other characterized plant GESs (Figure 4d). Additionally, we analyzed CfG(P)Pase alongside putative orthologs from both geraniol-producing and non-geraniol-producing plants to gain insights into their evolutionary divergence and potential functional conservation (Figure 4e). GESs belong to the TPS-b subfamily of TPS, which is primarily associated with monoterpene biosynthesis. Candidates of TPS-b subfamily harbor DDxxD and NSE/DTE motifs essential for the cofactor (Mg^2+^ or Mn^2+)^ binding to catalyze the synthesis of monoterpenes and N-terminal RRX8W domain involved in the cyclization of monoterpenes (Iijima et al., 2004) and (Zhao et al., 2020). The RRX8W motif, which is often, but not always, found in the N terminus of mature monoterpene synthases is not found in GES, consistent with the hypothesis that is involved in the synthesis of cyclic terpenes (Williams et al., 1998). Moreover, multiple sequence alignment (MSA) of CfTPS1 and several other known geraniol synthases (GES) revealed that CfTPS1 shares core functional motifs (Figure S6). Specific residues, such as those involved in the binding of divalent metal ions (e.g., Mg^2+^), are conserved and typically part of the DDxxD motif which is essential for the catalytic function of terpene synthases. The phylogenetic tree for GESs indicates that CfGES is part of a different evolutionary clade similar to GES from rice, which suggest functional or evolutionary divergence from the typical GESs of dicot plant species (Figure 4d). Similarly sequence analysis of CfPPase1 showed that it belongs to superfamily of pyrophosphatases which can hydrolyze inorganic pyrophosphate (PPi) to two molecules of orthophosphates (Pi) (Figure S7). Further phylogenetic analysis of CfPPase1 along with CfPPase2 and other predicted pyrophosphatases from plants indicated that CfPPase1 groups distinctly from other enzymes, potentially indicating it has unique functional or evolutionary characteristics compared to the more distantly related phosphatases (Figure 4e). As both CfGES and CfG(P)Pase contributed for geraniol formation in lemongrass, we searched the existence of their orthologs in other plants known to produce geraniol. Interestingly, most of the geraniol-producing plants such as *C. roseus*, consisted of both GES and PPase candidates indicating the potential presence of both canonical TPS-dependent and non-canonical PPase-dependent pathways for geraniol production.

In conclusion, our study uncovers dual routes for geraniol biosynthesis in lemongrass, mediated by *CfGES* and *CfG(P)Pase* (Figure 4f). To our knowledge, this is the first report of a plant pyrophosphatase that utilizes GPP and GP to produce geraniol. These findings expand our understanding of monoterpene biosynthesis in plants and highlight the existence of both TPS-dependent and TPS-independent pathways for geraniol formation not only in lemongrass but also in other geraniol producing plants. This knowledge has implications for crop improvement of aromatic grasses and metabolic engineering for production of geraniol-derived essential oils.

## Material and Methods

### Plant material and tissue collection

*Cymbopogon flexuosus cv*. Krishna (National Gene Bank, CSIR-CIMAP, Lucknow, India) was used for all experimental procedures. For crude protein assays and volatile compound extraction, leaves from mature plants grown under field conditions was utilized. To determine stage-specific gene expression analysis, leaves from various developmental phases (1 month, 2 months, 3 months, and the harvesting stage) were collected and utilized for RNA isolation and gas chromatography analysis. For subcellular fractionation, young leaves from plants grown in natural field conditions was collected and utilized.

### Structure modelling and molecular docking

For molecular docking analysis we generated three-dimensional models of CfGESTPS1 and CfG(P)Pase1 using AlphaFold 3, based on their respective FASTA sequences. The first model generated for each protein, selected based on its confidence score and structural accuracy, was then subjected to energy minimization using the Molecular Mechanics Toolkit within UCSF Chimera 1.15 (Pettersen et al., 2004). Following minimization, the protein structures were prepared for docking in Auto Dock Tool 1.5.7 (Rizvi et al., 2013). This preparation involved adding polar hydrogens and Kollman charges to the proteins. Concurrently, three-dimensional conformer files for the ligands—Geranyl pyrophosphate/diphosphate (GPP), Geranyl phosphate (GP), Geraniol, and Farnesyl pyrophosphate/diphosphate (FPP)—were retrieved from PubChem and converted into docking-compatible formats using PyMol. These ligands were then prepared in Auto Dock Tools 1.5.7 by adding Gasteiger charges, merging non-polar hydrogens, and defining their torsional degrees of freedom. To define the probable binding sites on the target proteins, structural homologs were identified from the Protein Data Bank (PDB). Specifically, PDB entries 3q4w, 5ucq, and 5cux served as references for CfG(P)Pase1, while 2ong and 3n0g were used for CfGES. Molecular docking was subsequently performed using Auto Dock Vina 1.2.5. For CfG(P)Pase1 with GPP, FPP, and GP, and for CfGES with GPP and Geraniol, the docking calculations were carried out within a grid box of 18.375 x 18.375 x 18.375 Å with a grid spacing of 0.375 Å. A smaller grid box, measuring 10.875 x 10.875 x 10.875 Å with the same spacing, was used for the docking of CfG(P)Pase1 with Geraniol. The resulting docked complexes were evaluated, and the complex exhibiting the most favourable conformation and the lowest binding energy was selected as the representative docked pose for each protein-ligand pair.

### RNA isolation, cDNA synthesis and qRT-PCR analysis

Total RNA was extracted from leaves of lemongrass, and lemon balm, and petals of rose following the protocol of (Deepa et al., 2014) with slight modifications. Briefly, 50 mg of leaf tissue was flash-frozen in liquid nitrogen and finely ground into a powder, followed by the addition of 2% polyvinylpyrrolidone (PVP) in powdered form to minimize polyphenolic interference. The ground tissue was then mixed with 1 mL of extraction buffer, vortexed thoroughly, and subjected to sequential acid phenol:chloroform extraction to efficiently remove polyphenols and proteins. Further purification was performed using sodium acetate to eliminate polysaccharide contaminants. The concentration and purity of the extracted RNA were determined spectrophotometrically by measuring absorbance at A260 and evaluating the A260/280 ratio. cDNA was synthesized from the isolated RNA and subsequently used for qRT-PCR analysis as described previously (Singh et al., 2015; Singh et al., 2017) *EF1α* served as the internal reference for cDNA normalization. For qRT-PCR, gene-specific primers were designed outside the regions used for cloning into pTRV2 (Table S2). The reaction was carried out using a 2X SYBR Green mix (Thermo Scientific, USA) in a StepOne Real-Time PCR System (Applied Biosystems, USA). The qRT-PCR cycling conditions were set as follows: initial denaturation at 94°C for 10 min, followed by 40 cycles of 94°C for 15 s and 60°C for 45 s. Relative gene expression levels were quantified using the comparative cycle threshold (Ct) method to assess fold-change differences.

### Protein expression and purification

The coding sequence of *CfTPS1* was cloned into pDEST527 bacterial expression vector through gateway cloning system and that of *CfPPase1* was cloned into the *Nde*I*/Eco*RI restriction sites of the pET28a bacterial expression vector, ensuring an in-frame fusion with a C-terminal 6X-His-tag (Novagen, http://www.emdbiosciences.com). This resulted in the recombinant constructs pDEST527*::CfTPS1* and pET28a*::CfPPase1*. For recombinant protein expression, these constructs, along with empty pDEST527 and pET28a vector control, were introduced into *E. coli* BL21 Rosetta-2 cells. A single transformed colony was cultured overnight in 5 mL of LB medium containing kanamycin (50 mg/L) and chloramphenicol (34 mg/L) at 37°C with shaking at 180 rpm. The overnight culture (1 mL) was then inoculated into 500 mL of fresh LB medium with the same antibiotics and grown at 37°C until the optical density at 600 nm (OD600) reached 0.4. At this stage, protein expression was induced by adding 0.4 mM isopropyl-β-D-thiogalactopyranoside (IPTG), followed by incubation at 16°C with shaking at 180 rpm for 16 hours. Recombinant proteins were purified using nickel-nitrilotriacetic acid (Ni-NTA) agarose beads (Qiagen, http://www.qiagen.com) according to the manufacturer’s protocol. The protein concentration was determined using the Bradford assay (Bradford, 1976), and purity was assessed by SDS-PAGE analysis (Figure. S9).

### Enzyme assay of recombinant CfGES and CfG(P)Pase

Enzyme assays were conducted utilizing purified recombinant proteins, CfTPS1 and CfPPase1. The assay buffer was composed of 10 mM MOPSO at pH 7.5, 1 mM MgCl_2_, 0.1 mM mnCL_2_ and 5 mM DTT. Each purified enzyme (CfTPS1 and CfPPase1) was incubated at a concentration of 5-10 µg with 50 μM of the corresponding substrate in a 500 μL reaction mixture, which was subsequently overlaid with 500 μL of n-hexane. The reaction was maintained at 30°C for a duration of 2 hours. After the incubation period, the upper hexane layer, which contained the volatile products, was carefully transferred into a separate vial. The hexane extract was then concentrated and preserved at −20°C for future analysis. For gas chromatography (GC) analysis, 10 μL of the concentrated extract was injected, while for gas chromatography-mass spectrometry (GC-MS), 3 μL was utilized to identify and quantify the volatile compounds.

### Generation of silencing and overexpression constructs

The vectors pTRV1 and pTRV2 used for creating VIGS constructs were obtained from The Arabidopsis Information Resource (TAIR) in the United States. Fragments measuring 551 bp, 523 bp, and 502 bp corresponding to *CfPDS, CfGES*, and *CfPPase1*, respectively, were amplified from leaf cDNA through PCR, employing gene-specific primers that included an *EcoR*I restriction site at the 5′ end for *CfGES*, and *CfPPase1*, and a *Bam*HI site for *CfPDS* (refer Table S2). The resultant PCR products were purified and subsequently sub-cloned into the pJET1.2 vector, and sequences were verified through nucleotide sequencing. The confirmed fragments were then inserted into the pTRV2 vector, which was digested with the appropriate restriction enzymes. The constructs pTRV2::*CfPDS*, pTRV2::*CfGES*, and pTRV2::*CfPPase1* were validated via restriction digestion. For generating overexpression constructs, the open reading frames of *CfGES* and *CfPPase1* were PCR amplified using cDNA derived from lemongrass leaves, utilizing specific forward and reverse primers (refer Table S2). The resulting amplicons were cloned into the pJET1.2 cloning vector for sequence verification, and subsequently sub-cloned into the *Nde*I and *Eco*RI sites of the pRI101-AN binary vector, under the regulation of the 35S promoter from Cauliflower mosaic virus (CaMV), resulting in the pRI::*CfGES* and pRI::*CfPPase1* constructs (illustrated in Figure. S10). The generated VIGS and overexpression constructs were mobilized into *Agrobacterium tumefaciens* GV3101 competent cells using a freeze-thaw method.

### Silencing of *CfGES* and *CfPPase1* in lemongrass through VIGS

Overnight cultures of *Agrobacteria* containing pTRV1 and various constructs, including pTRV2*::CfPDS*, pTRV2*::CfGES*, pTRV2*::CfPPase1*, and pTRV2 (EV), were resuspended in a MES buffer solution composed of MES, MgCl_2_, and Acetosyringone. These cultures were mixed in a 1:1 ratio and incubated on a shaker for three hours. Subsequently, approximately 100 µl of the agro-suspension was injected into the tiller-base region of uprooted seedlings to facilitate infection. The infected seedlings were then replanted into pots filled with soilrite, placed in a box covered with perforated cling film to maintain humidity, and kept in the dark for 48 hours. After this period, the pots were transferred to a growth room maintained at 22 °C with a 16-hour light and 8-hour dark cycle. Four weeks post-infection, leaves from plants infected with pTRV2*::CfPDS*, pTRV2*::CfGES*, pTRV2*::CfPPase1*, and pTRV2 were harvested, at which point the albino phenotype was observed in the newly emerging leaves of the pTRV2*::CfPDS* infected plants. The harvested tissues were subsequently divided into two portions: one for immediate volatile extraction and the other stored at −80°C for future RNA extraction.

### Transient expression of *CfADH1* and *CfAKR2b* in lemon balm and Rose

Transient expression was conducted using lemon balm plants at the 6-8 leaf stage. *Agrobacterium* strains containing p19 (an RNA silencing suppressor) along with pRI*::CfGES*, pRI*::CfPPase1*, or a control pRI construct were combined in a 1:1 ratio (OD_600_=0.6) and incubated in a shaker at 28°C for 3 hours. Agroinfiltration was executed in a vacuum desiccator by submerging the upper portion of the lemon balm seedlings, including all leaves, into the Agrobacterium solution in an inverted position, followed by the application of vacuum for 10-15 minutes. Post-infiltration, the plants were placed in a box covered with perforated cling film to retain humidity and kept in the dark for 48 hours. Subsequently, the plants were taken out of the box and transferred to a growth room maintained at 22°C with a 16-8-hour light cycle for 4 days. Leaf samples (the top four) were then collected for immediate volatile and RNA extraction. Similarly, transient overexpression in rose petals was done using a needleless syringe by gently pressing the syringe against the petal surface and infiltrating the Agrobacteria solution until the petal area became visibly saturated. The infiltrated petals were kept in a humid chamber in the dark for 48 h and then transferred to normal growth conditions for 24 h before sample collection for volatile and RNA extraction.

### Extraction and quantification of volatiles

For metabolite extraction, 50 mg of lemongrass leaves and 100 mg of lemon balm leaves and rose petals were used. The weighed tissue was placed in an extraction vial with 1 mL of hexane containing the internal standard (*E,E*)-farnesol. The mixture was incubated at 28°C overnight to facilitate metabolite extraction. After incubation, the samples were centrifuged at 5000 *g* for 5 min, and the supernatant was carefully collected. To remove residual water, the hexane extract was treated with anhydrous Na_2_SO_4_, and activated charcoal was added to eliminate chlorophyll and other pigments. The extract was then filtered through a 0.22 µm syringe filter to remove any remaining particulates and stored at −20°C until further analysis. In all experiments, volatile terpene compounds were detected and identified using gas chromatography (GC) and/or gas chromatography-mass spectrometry (GC-MS). For precise quantification, calibration curves were established using authentic standards of the target volatile compounds, and their concentrations in the samples were determined via GC-MS analysis. Additionally, the concentration of volatiles was further estimated using the Relative Response Factor (RRF) method in GC, with *E,E*-farnesol serving as the internal standard to ensure accuracy and reproducibility in quantification.

### Analysis of subcellular localization

The subcellular localization of *CfGES* and *CfG(P)Pase* was investigated by generating YFP fusion proteins using the pGWB441 vector. The coding sequences of *CfGES* and *CfG(P)Pase* were cloned into pGWB441 to express them as C-terminal eYFP-tagged proteins, resulting in the constructs pGWB441*::CfGES-YFP* and pGWB441*::CfG(P)Pase-YFP*. These constructs were introduced into *Agrobacterium tumefaciens* strain GV3101 via the freeze-thaw transformation method. For transient expression in *Nicotiana benthamiana* leaves, *Agrobacterium* cultures were resuspended in infiltration buffer (10 mM MES, pH 5.6, 10 mM MgCl_2_, and 200 μM acetosyringone). Prior to infiltration, *Agrobacterium* carrying the overexpression construct or empty vector (EV) was mixed with an *Agrobacterium* culture containing the *p19* suppressor of RNA silencing in a 1:1 ratio. Agroinfiltration was performed on 5- to 6-week-old *N. benthamiana* plants. After 36 to 48 hours post-infiltration, leaf sections were examined under a Carl Zeiss LSM880 laser scanning confocal microscope using a 63× oil immersion objective (numerical aperture 1.4). YFP fluorescence was excited at 514 nm and detected in the 525–562 nm range, while chlorophyll autofluorescence was excited at 637 nm and detected in the 660–700 nm range. Fluorescent images were captured to determine the precise subcellular localization of the fusion proteins.

### Separation of subcellular fractions

Subcellular fractionation of lemongrass leaves was performed following the protocol described by (An et al., 2021), with slight modifications. Young leaves from field-grown plants were cut into 1–3 cm segments, and 7 g of tissue was homogenized in 100 mL of grinding buffer (GB) containing 50 mM HEPES/KOH (pH 7.9), 0.33 M sorbitol, 1 mM MgCl_2_, 1 mM MnCl_2_, 2 mM EDTA (pH 8.0), 5 mM sodium ascorbate, and 0.1% bovine serum albumin (BSA). The homogenization was carried out using a kitchen blender with five pulses of 3 seconds each. The homogenized mixture was filtered through Miracloth into 50 mL tubes and centrifuged at 2,000 g for 2 minutes. The resulting supernatant, representing the cytosolic fraction, was collected, while the pellet containing chloroplasts was retained for further purification. The pellet was resuspended in 500 µL of resuspension buffer (RB) composed of 50 mM HEPES-KOH (pH 7.9), 0.33 M sorbitol, 0.8 mM CaCl_2_, and 5 mM sodium ascorbate. This suspension was carefully layered onto a Percoll density gradient (40% Percoll on top and 90% Percoll at the bottom) and centrifuged at 2,000 g for 10 minutes. The intact chloroplasts, appearing in the second layer, were collected, transferred to a fresh tube, resuspended in 3 mL of RB, and subjected to centrifugation at 400 g for 3 minutes. The purified chloroplasts were then resuspended in an assay buffer containing 15 mM Tris-HCl (pH 7.5), 1 mM DTT, 5 mM MgCl_2_, and 10% glycerol. To confirm the purity of the cytosolic fraction, the activity of the cytosolic marker enzyme alcohol dehydrogenase was measured (Denyer and Smith, 1988). The integrity of the isolated chloroplasts was verified using confocal microscopy. Chloroplast images were captured using a Carl Zeiss LSM880 laser scanning confocal microscope with a 63× oil immersion objective (numerical aperture 1.4). Chlorophyll autofluorescence was excited at 634 nm and detected in the 571–696 nm range. Following integrity verification, chloroplasts were lysed by sonication using three pulses of 35 seconds each. To eliminate background volatiles, all subcellular fractions were desalted before further analysis.

### Phylogenetic analysis

For phylogenetic analysis, amino acid sequences of characterized GESs along with putative pyrophosphatase sequences from different plant species were retrieved from the National Center for Biotechnology Information (NCBI) database (www.ncbi.nlm.nih.gov). The phylogenetic tree was generated using the neighbor-joining method in MEGA11 software (Tamura et al., 2021). Multiple sequence alignment was conducted using ClustalW (Thompson et al., 1994) with default parameters via the online tool (https://www.genome.jp/tools-bin/clustalw). The accession numbers of the sequences utilized for tree construction are listed in Table S3.

## Accession numbers

CfGES (PV167775); CfG(P)Ppase (PV167776).

## Acknowledgements

This research was supported by the MLP0003 and HCP0007 projects of Council of Scientific and Industrial Research. P.G., and A.S., & S.M., are the recipients of senior research fellowship from Department of Biotechnology (DBT), and University Grants Commission (UGC), respectively. The authors are thankful to Drs. Sumit Ghosh for sparing pGWB441 vector, Manish Tiwari for providing *Nicotiana benthamiana* plants, and Rajendra Patel for help with confocal analysis. The authors also express their sincere gratitude to the Director, CSIR-CIMAP for support throughout the study. Institutional communication number for this article is CIMAP/Pub/2025/143. R.N. Lab work was supported by core research grant to IISER-Thiruvananthapuram from Ministry of Education (MoE), Government of India. R.N. thanks MoE STARS (STARS/APR2019/BS/729/FS) and SERB-CRG (CRG/2023/001211) for the financial support. R.N. and Y.M. would like to thank the Padmanabha cluster at CHPC, IISER-TVM and also like to thank Professors E.D. Jemmis, V. Ramakrishnan and J.N. Moorthy, past and present directors for their unstinted support. Authors declare no conflict of interest.

## Author contributions

DAN planned and designed the research. PG, AS, SM, and YM performed experiments, conducted fieldwork, and analysed data. DAN, RN, and PG analysed and interpreted the data. DAN and PG and wrote the manuscript. DAN, PG, YM and RN worked on first draft.

## Data availability

The authors confirm that the data supporting the findings of this study are available within article in Supporting Information.

