## Supplemental Information for "A dual-localized geraniol synthase and a previously unreported cytosolic geranyl pyrophosphatase contribute to geraniol formation in lemongrass"

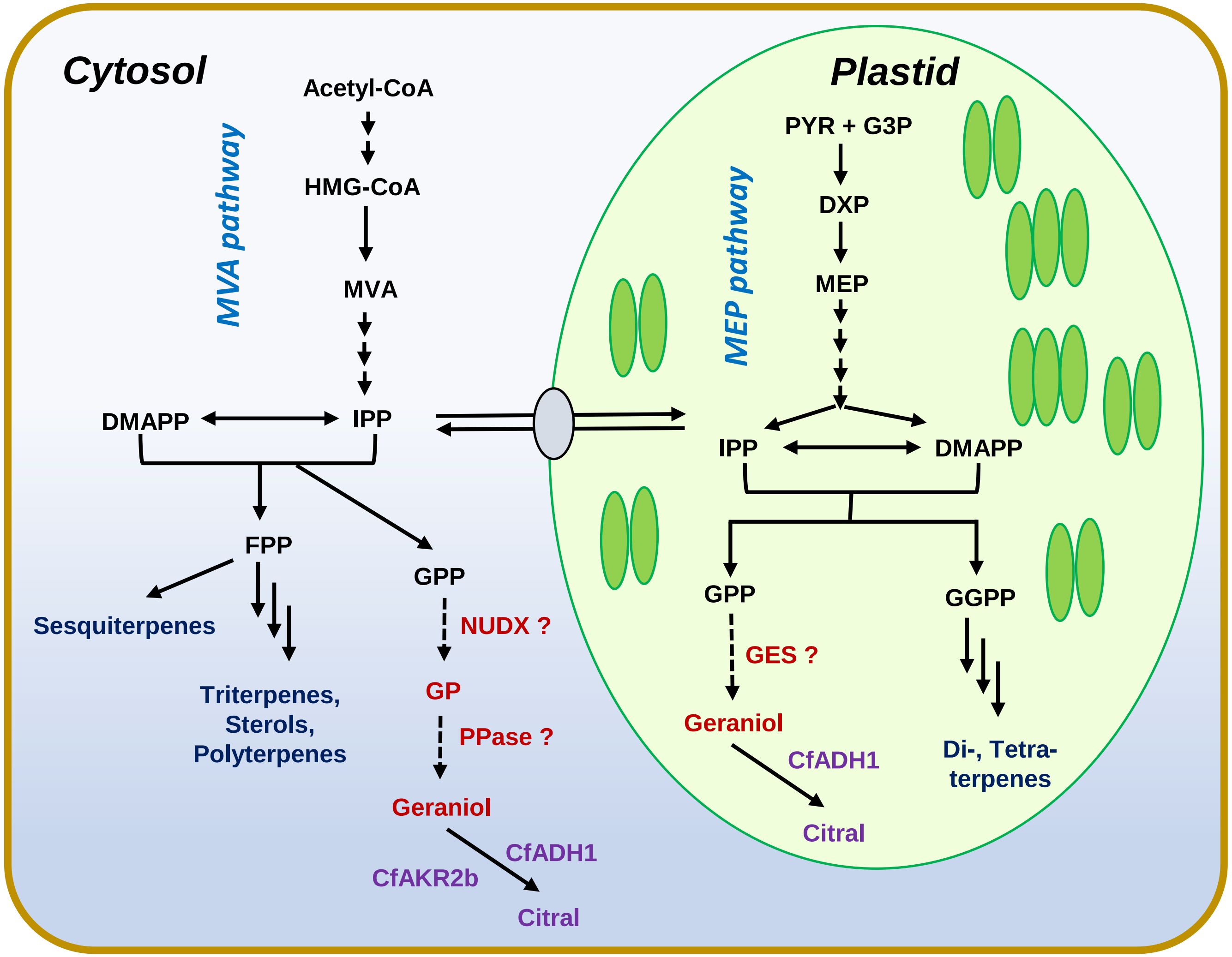


**Figure S1. Possible routes of geraniol biosynthesis in lemongrass.** Dotted arrows indicate putative steps. Question mark indicates possible enzyme involved in the pathway. Already characterized enzymes in lemongrass are in purple color. Abbreviations: CfADH1, *Cymbopogon flexuosus* alcohol dehydrogenase 1; CfAKR2b, *C. flexuosus* aldoketo reductase 2b; DMAPP, dimethylallyl diphosphate; DXP, 1-deoxy-D-xylulose 5-phosphate; FPP, farnesyl diphosphate; IPP, isopentenyl diphosphate; G3P, glyceraldehyde 3-phosphate; GPP, geranyl diphosphate; GGPP, geranylgeranyl diphosphate; GES, geraniol synthase; HMG-CoA, 3-hydroxyl-3-methylglutaryl CoA; MEP, methylerythritol phosphate; MVA, mevalonic acid; PPase, pyrophosphatas; PYR, pyruvate.

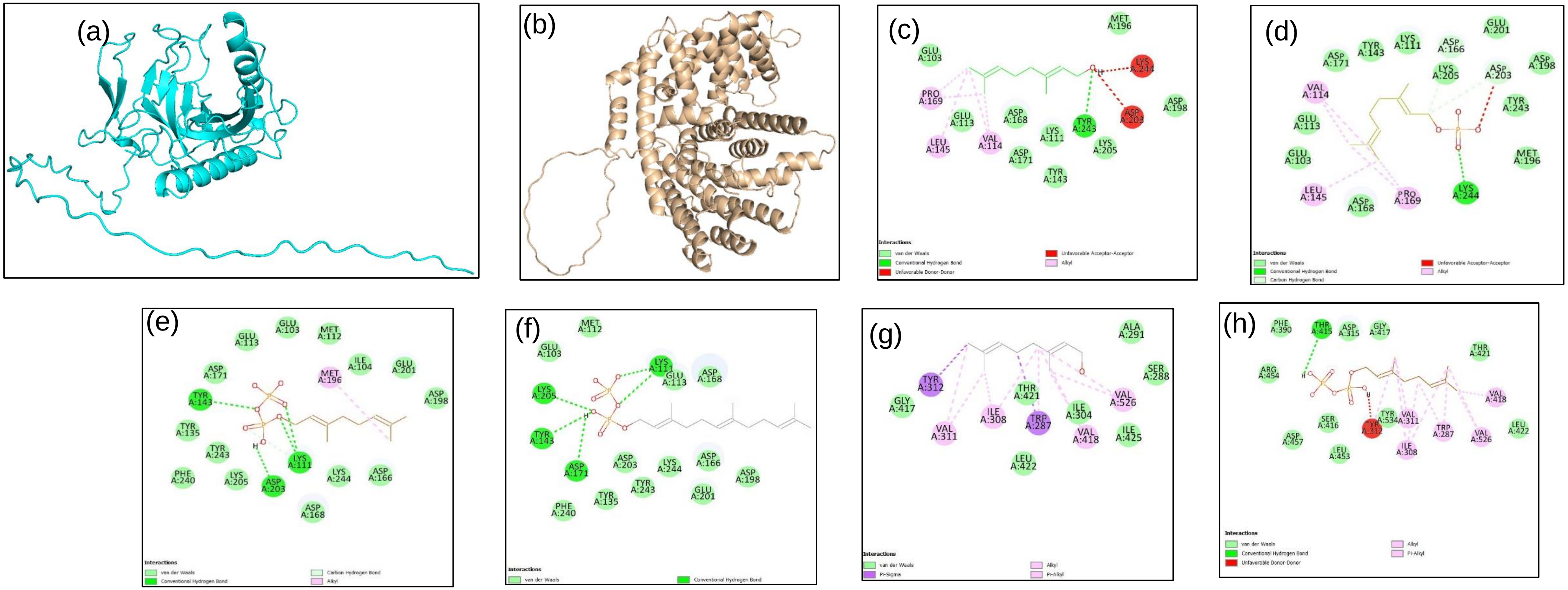


**Table 1. Details of docking results.**

| Protein | Ligand | Binding Affinities (kcal/mol) | No. of H-Bonds | Amino Acids involved in H-Bonds |
|---|---|---|---|---|
| **CfPPase1** | Geraniol | -4.6 | 1 | Tyr243 |
| | GP | -5.0 | 1 | Lys244 |
| | GPP | -5.8 | 4 | Lys111, Tyr143, Asp203 |
| | FPP | -5.7 | 5 | Lys111, Tyr143, Asp171, Lys205 |
| **CfTPS1** | Geraniol | -6.4 | 0 | |
| | GPP | -7.9 | 1 | Thr415 |

**Figure S2. Modelling and Docking analysis:** Alpha Fold 3 modelled structures of CfPPase1 (a) and CfTPS1 (b). 2D representation of ligand-protein interaction of Geraniol docked to CfPPase1 (c ) GP docked to CfPPase1 (d) GPP docked to CfPPase1 (e) FPP docked to CfPPase1 (f) Geraniol docked to CfTPS1 (g) GPP docked to CfTPS1 (h).

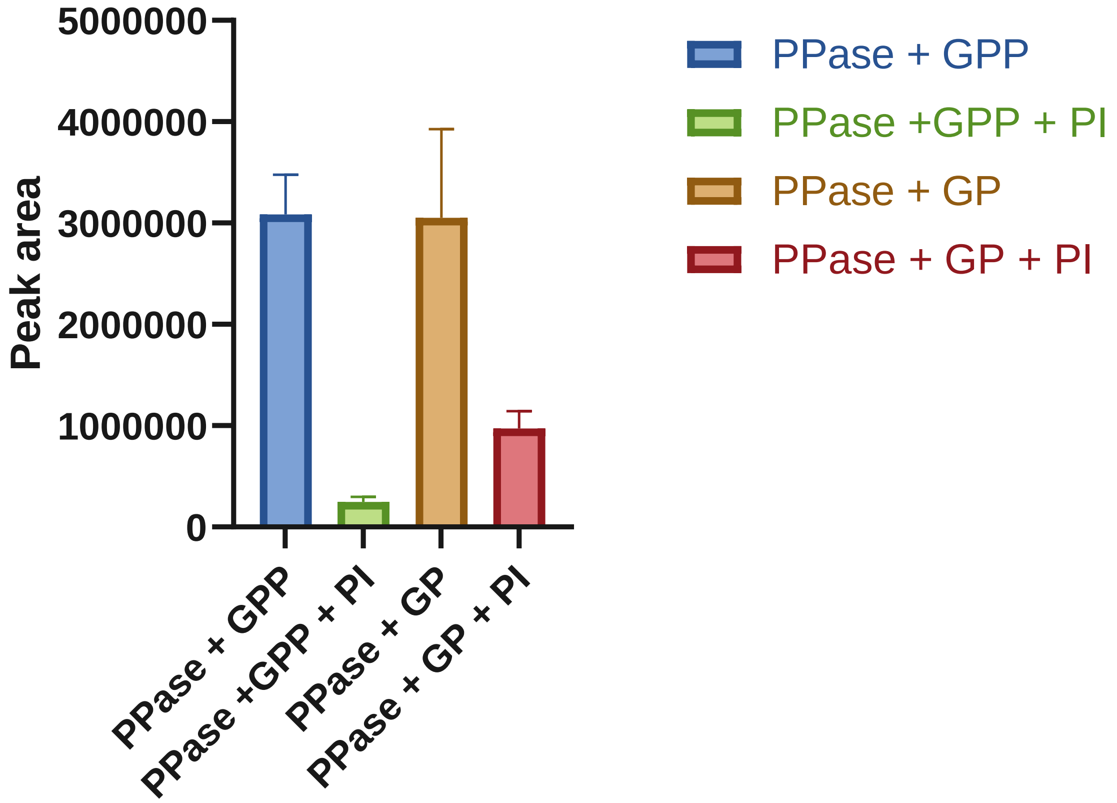


**Figure S3. Recombinant CfG(P)Pase enzyme assay in the presence of phosphatase inhibitor (PhosSTOP, Sigma-Aldrich) using GPP and GP as substrates.**

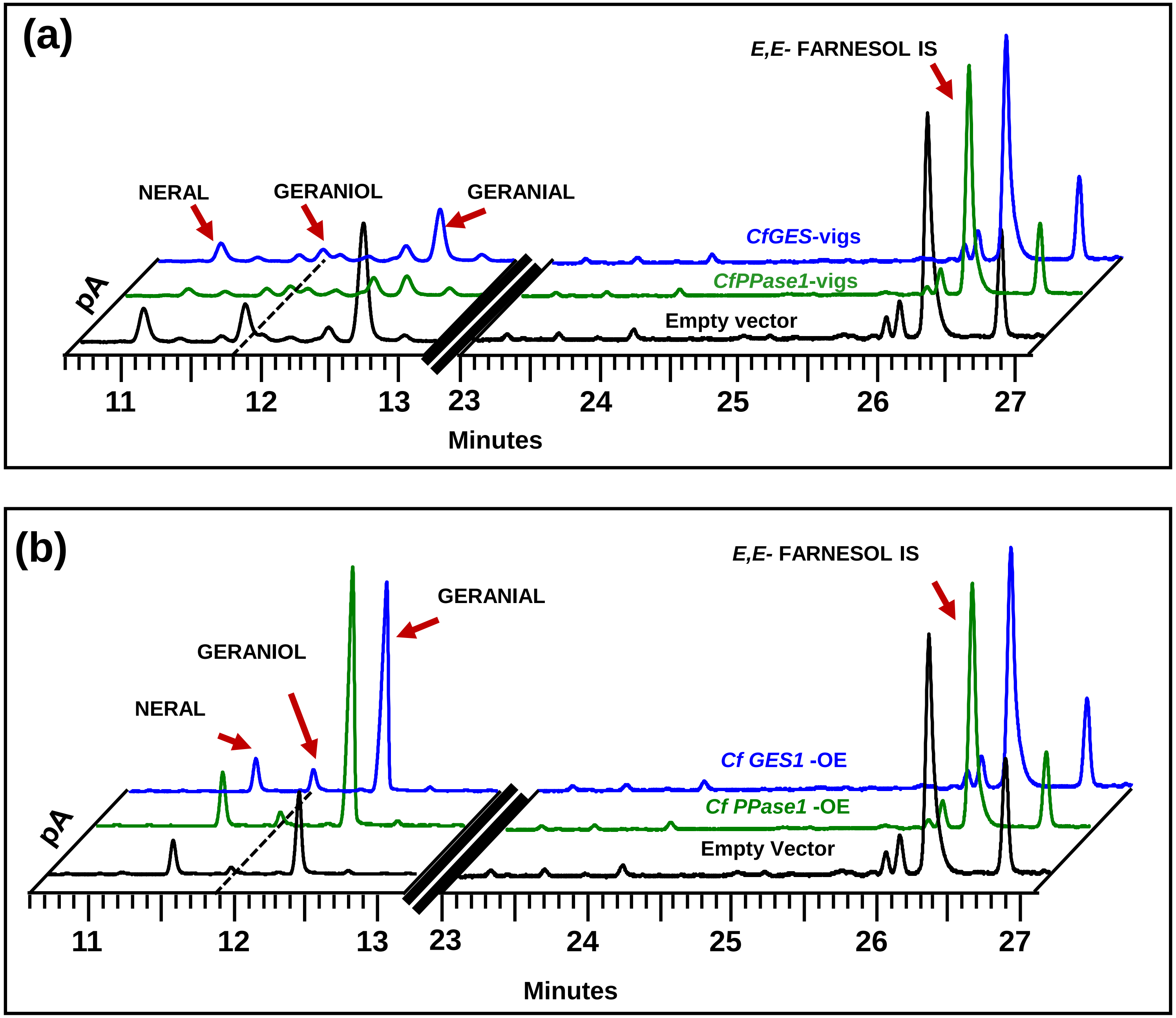


**Figure S4:** (a) Representative chromatograms from EV control, *CfGES*-vigs, and *CfPPase1*-vigs showing geraniol and citral peaks. (b) Representative chromatograms from EV control, *CfGES*-OE, and *CfPPase*-OE showing geraniol and citral peaks. Volatiles were extracted using hexane containing internal standard (IS) E,E-farnesol, and subjected to gas chromatography (GC) analysis.

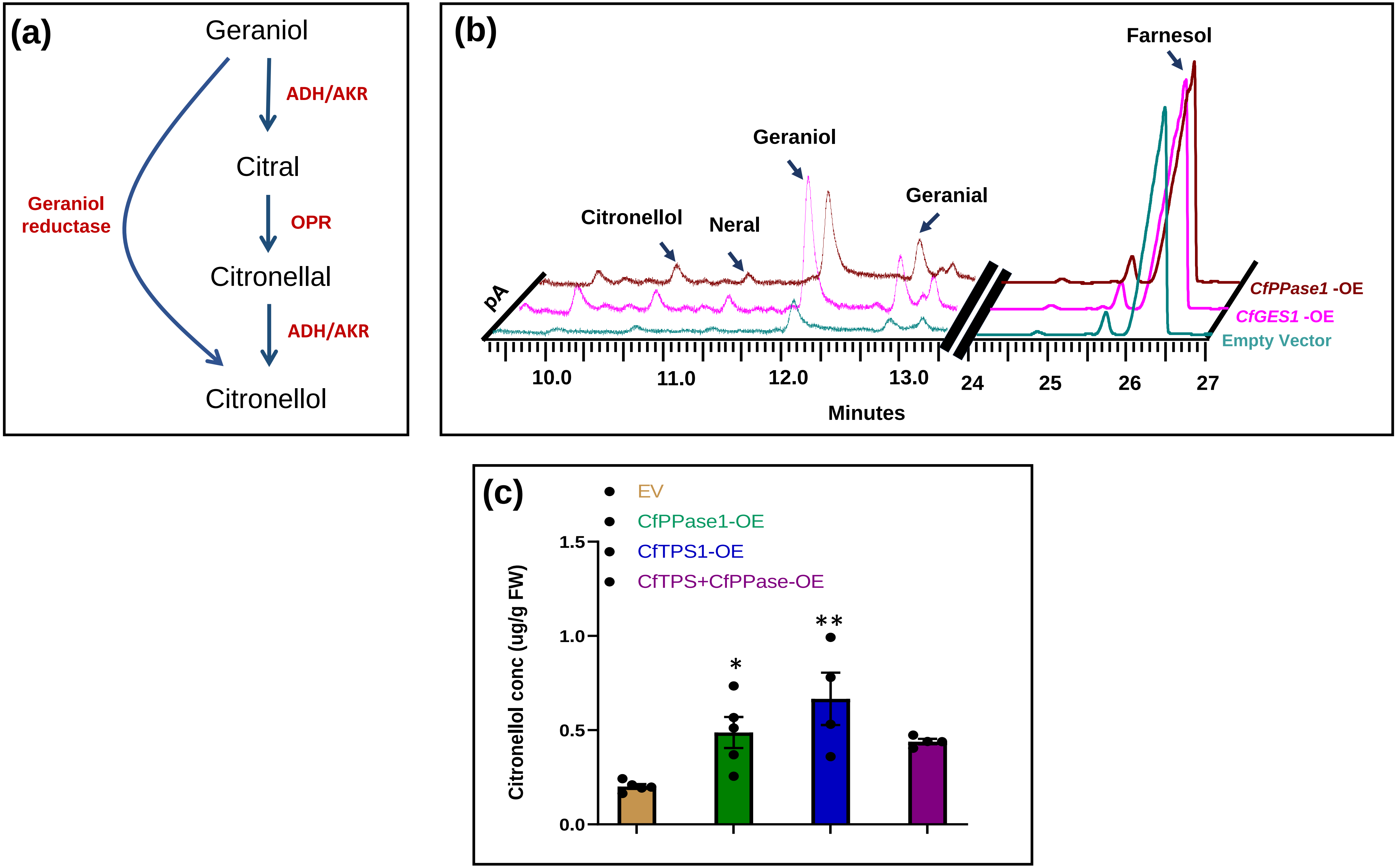


**Figure S5. (a)** Two different modes of citronellol biosynthesis in plants. First, geraniol can directly convert into citronellol as shown in Rose and Pelargonium (refs). Second is three step pathway where geraniol get converted into citral (refs( followed by citronellal and then citronellol (refs,PRISE). (b) Representative chromatograms from EV control, *CfGES*-OE, and *CfPPase1*-OE showing geraniol, citronellol and citral peaks. (c) Quantification of citronellol in EV control, *CfGES*-OE, and *CfPPase1*-OE tissues.

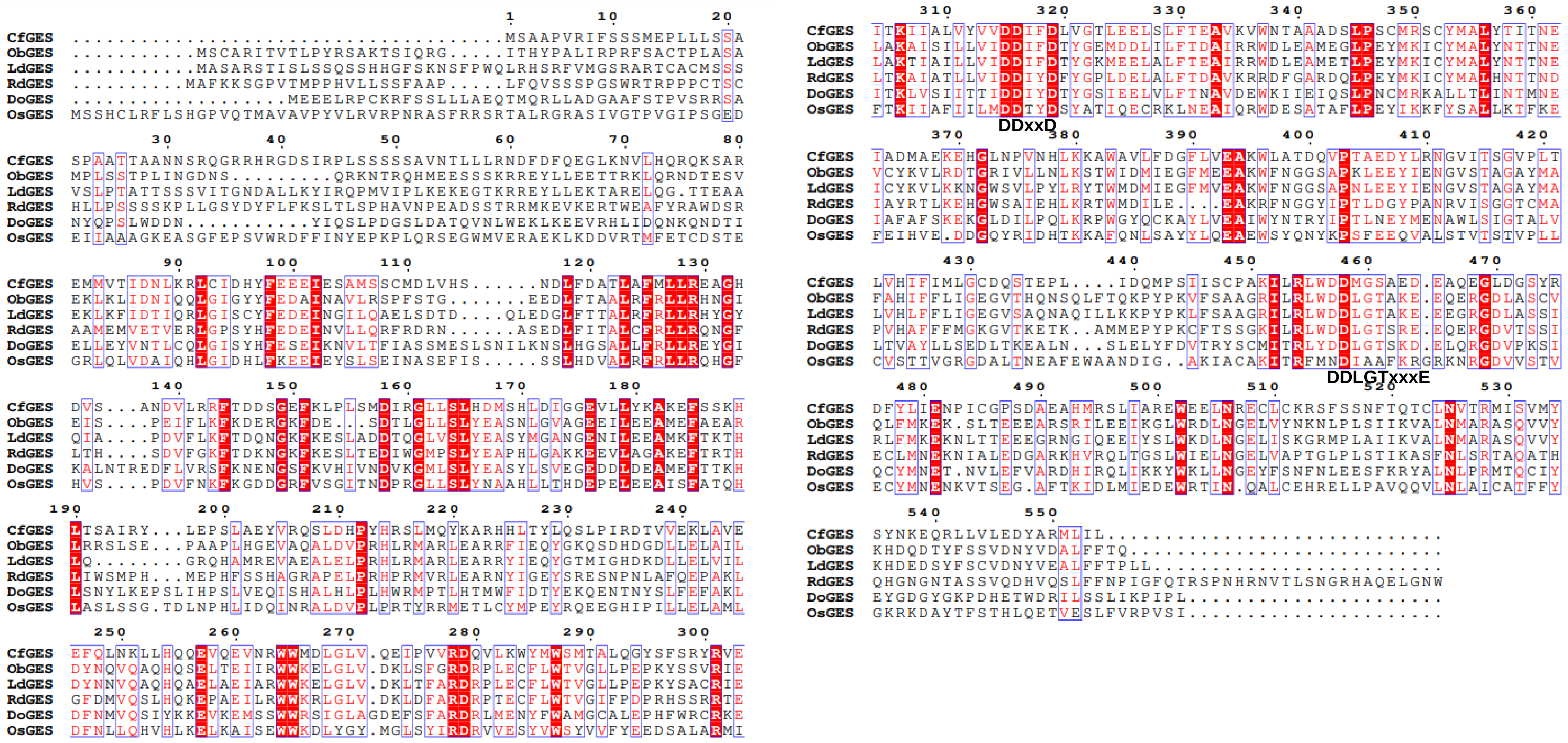


**Figure S6. Multiple sequence alignment of CfTPS1 with ObGES, LdGES, VoGES DoGES and OsTPS21.**

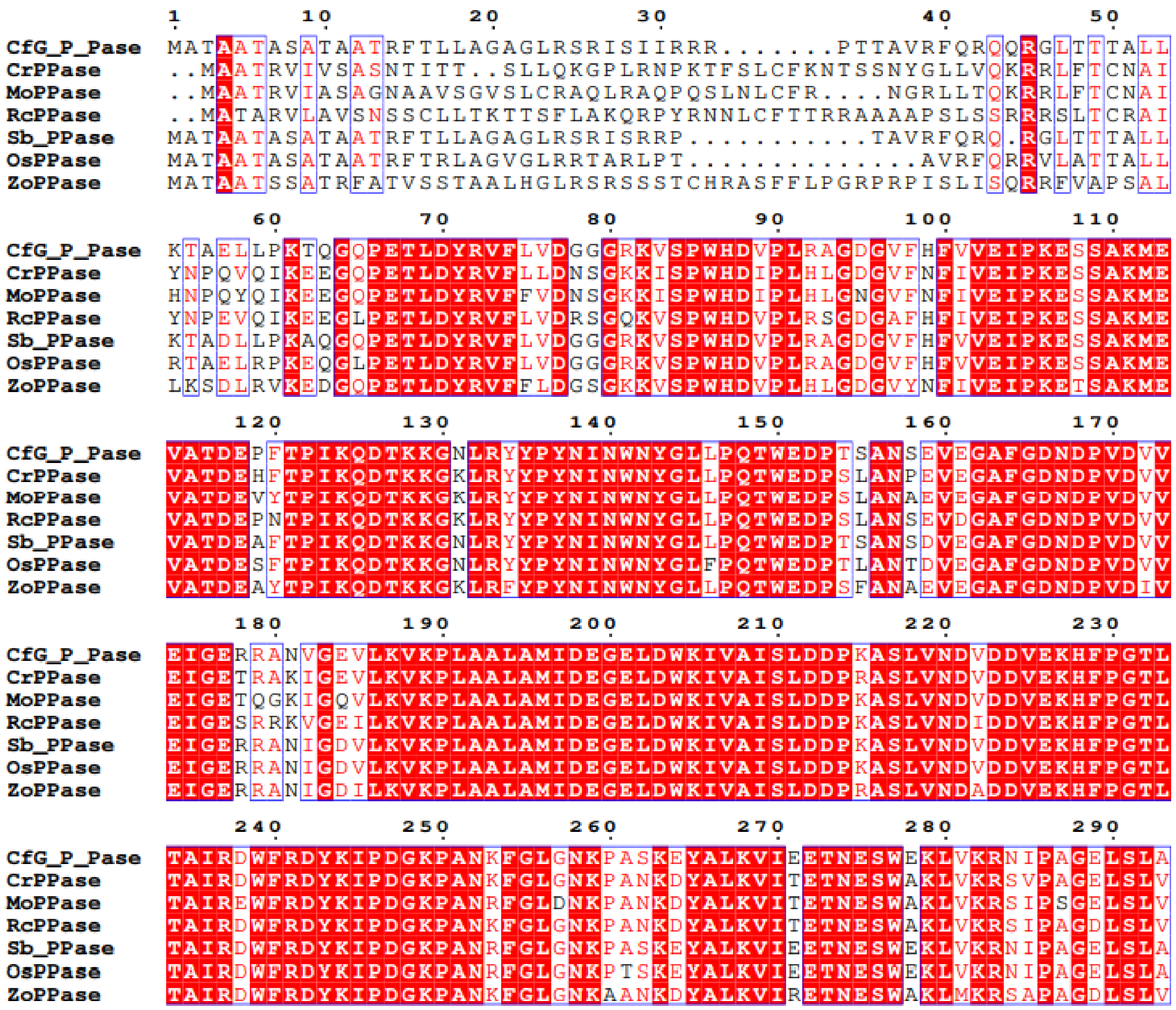


**Figure S7. Multiple sequence alignment of CfPPase with other putative plant pyrophosphatase.**

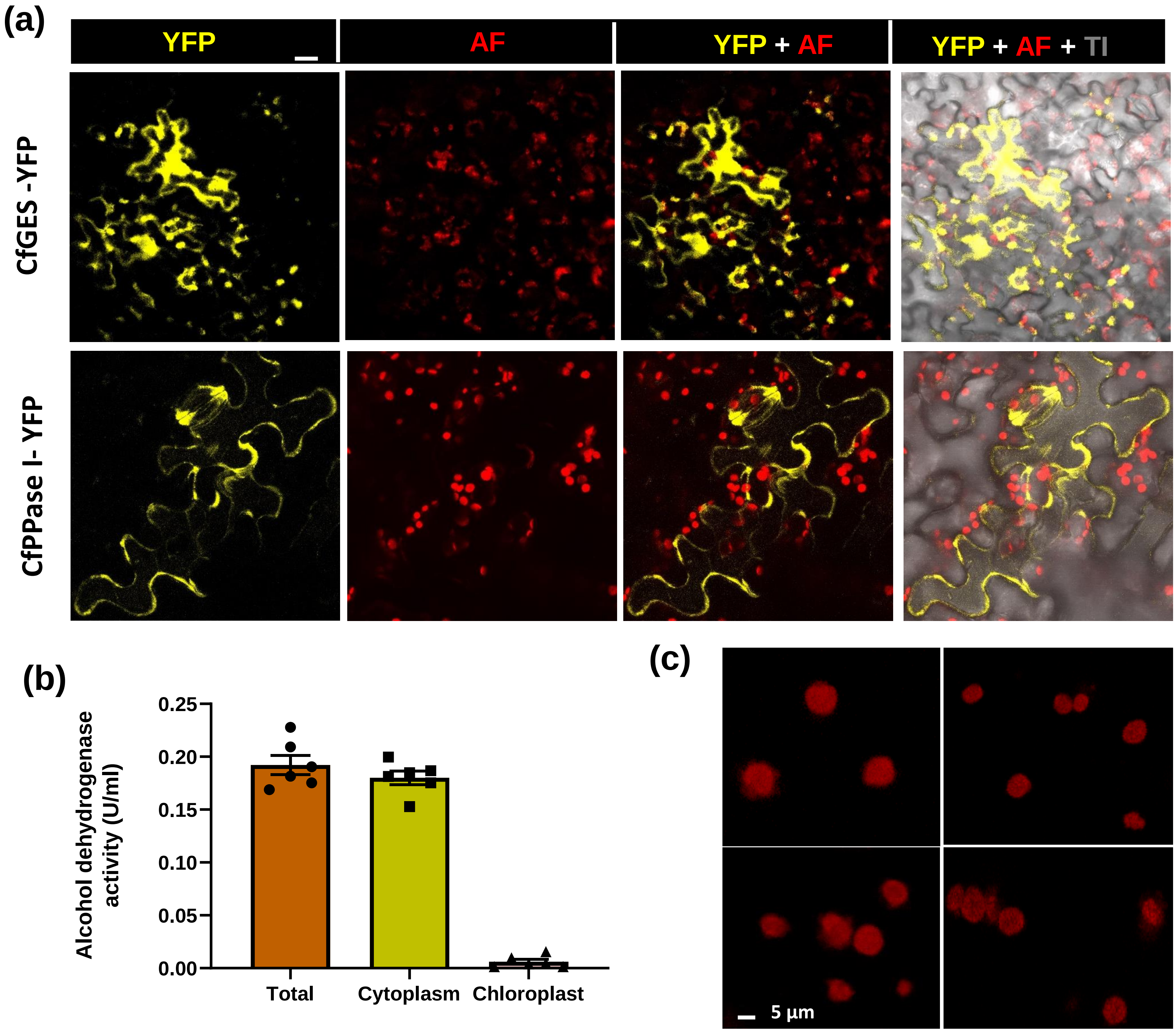


**Figure S8.** (a) Additional confocal images of CfGES-YFP and CfPPase1-YFP expressing cells. (b) Determination of purity of subcellular fractions using alcohol dehydrogenase assay. (c) Confocal images showing the integrity of chloroplasts in purified plastid fractions..

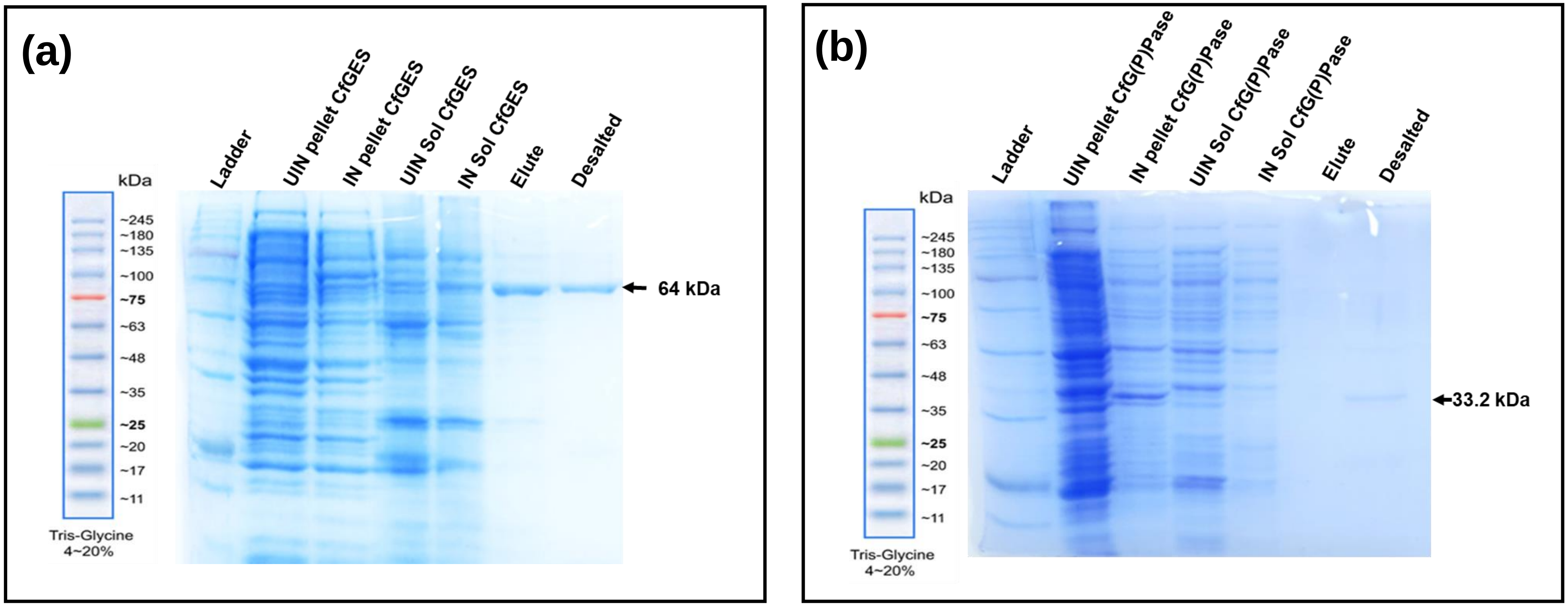


**Figure S9.** SDS-PAGE analysis of recombinant CfGES and CfG(P)Pase proteins expressed in *E. coli* Rosetta 2 cells. Recombinant proteins were purified from induced soluble fraction by Ni-NTA affinity chromatography. The elutes were desalted using assay buffer and stored at -80$^{o}$C for further use. Abbreviations: kDa, kilo Dalton. IN, induced; UIN, uninduced; Sol, soluble fraction;

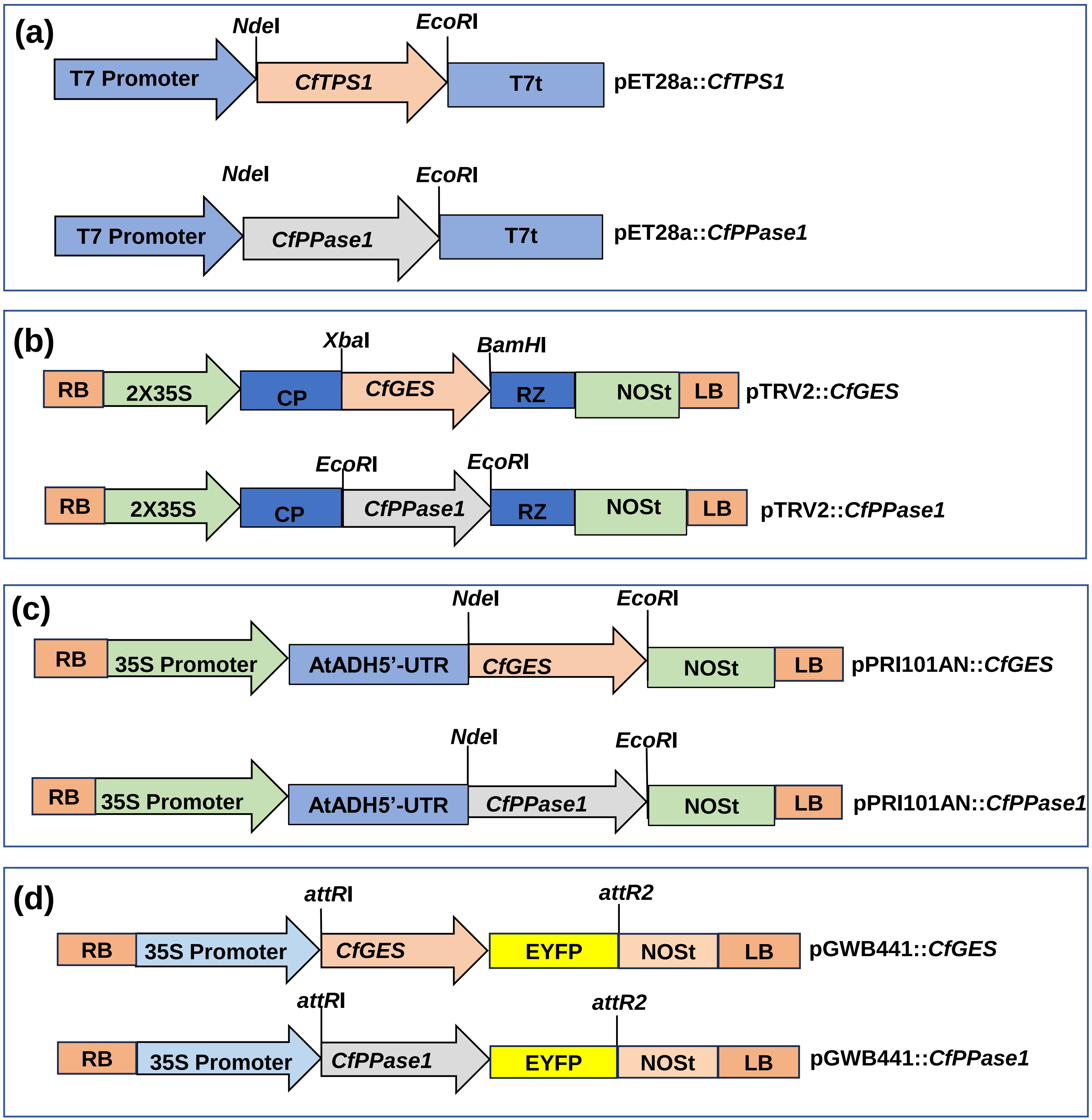


**Figure S10.** Schematic diagrams of constructs generated and used in this study. (a) pET28a-derived bacterial overexpression constructs. (b) pTRV2-derived VIGS constructs. (c) pRI101AN-derived plant overexpression constructs. (d) pGWB441-derived constructs used for subcellular localization. VIGS, virus induced gene silencing; AtADH, *Arabidopsis thaliana* alcohol dehydrogenase; NOST, Nopaline synthase terminator; UTR, untranslated region; CP, coat protein; RZ, self-cleaving ribozyme.

**Table S1.** *In silico* prediction of subcellular localization of CfGES and CfPPase1

| **Program** | **CfGES** | **CfPPase** |
|---|---|---|
| iPSORT<br>https://ipsort.hgc.jp/ | YES | YES |
| WOLFPSORT<br>https://wolfpsort.hgc.jp/ | chlo: 8 | chlo: 13 |
| Predotar<br>https://urgi.versailles.inra.fr/predotar/ | plastid | plastid |
| TargetP 2.0<br>https://services.healthtech.dtu.dk/services/TargetP-2.0/ | Other | Thylakoid luminal transfer peptide |
| Plant-mPLoc<br>http://www.csbio.sjtu.edu.cn/bioinf/plant-multi/ | Chloroplast. | Chloroplast.<br>Cytoplasm. |
| DeepLoc 2.0<br>https://services.healthtech.dtu.dk/services/DeepLoc-2.0/ | Plastid | Plastid |

**Table S2.** List of oligonucleotide primers used in this study.

| Sl No. | Gene | Sequences 5'---------------------------3' |
|---|---|---|
| 1 | CfTPS1_FL_F | GGTACCATGTCTGCTGCACCTGTA |
| 2 | CfTPS1_FL_R | GGATCCTCAAAGTATCAACATCC |
| 3 | CfPPase_FL_F | CATATGATGGCGACGGCGGCCACG |
| 4 | CfPPase_FL_R | GGATCCTTAGGCCAACGAGAGCTCTCC |
| 5 | CfTPS1_VIGS_F | TCTAGAATGTCTGCTGCACCTGTACGC |
| 6 | CfTPS1_VIGS_R | GGATCCCTCCAATGTCTAGGTGAGACATG |
| 7 | CfPPase_VIGS_F | GGATCCAAGAAGGGCAACCTTCGATAC |
| 8 | CfPPase_VIGS_R | GGATCCTTAGGCCAACGAGAGCTCTCC |
| 9 | CfPDS_F | GGATCCCACAATAAACTTTTTGGAAGCTGG |
| 10 | CfPDS_R | GGATCCCAGGAACACCCTGCTTTTTC |
| 11 | CfTPS1_RT_F | AGGCGGTCAAAGTGTGGAAT |
| 12 | CfTPS1_RT_R | GGTGTAAAGAGCCATGTAGCATGA |
| 13 | CfPPase_RT_F | CACCGACGAGCCCTTCAC |
| 14 | CfPPase_RT_R | CGGGTCCTCCCATGTTTG |
| 15 | CfPDS_RT_F | GCATTTTGATTGCTTTGAACAGA |
| 16 | CfPDS_RT_R | CCCTAGACCGAATGTGATCAACA |
| 17 | CfTPS1_GW_F | GGGGACAAGTTTGTACAAAAAAGCAGGCTT AATGTCTGCTGCACCTGTACG |
| 18 | CfTPS1_GW_R | GGGGACCACTTTGTACAAGAAAGCTGGGTT AAGTATCAACATCCTTGCATAGTCCTCAAG |
| 19 | CfPPase_GW_F | GGGGACAAGTTTGTACAAAAAAGCAGGCTT AATGGCGACGGCGGCCACG |
| 20 | CfPPase_GW_R | GGGGACCACTTTGTACAAGAAAGCTGGGTT TTAGGCCAACGAGAGCTCTCC |
| 21 | Mo_EF_1a_F | TGACAACGAAACGCAACACA |
| 22 | Mo_EF_1a_R | CATTGGGTACTTGACAGGCG |
| 23 | RdSAND_RT_F | GTGTTGAGGAGTTGCCTCTTG |
| 24 | RdSAND_RT_R | AACCTGTCGGGAGAATCTGTT |

**Table S3.** Accession numbers of characterized proteins used for phylogenetic analysis

| Protein name | Species | Accession number |
|---|---|---|
| CrGES | *Catharanthus roseus* | KF561459 |
| VoGES | *Valeriana officinalis* | KF951406 |
| ObGES | *Ocimum basilicum* | AY362553 |
| CtGES | *Cinnamomum tenuipilum* | AJ457070 |
| LdGES | *Lippia dulcis* | GU136162 |
| PfGES | *Perilla frutescens* | DQ234300 |
| VvGES | *Vitis vinifera* | NP_001267920 |
| DoGES1 | *Dendrobium officinale* | MT875214 |
| RdGES | *Rosa x damascena* | MN639696 |
| OsGES | *Oryza sativa L.* cv. Nipponbare | XP_015635171 |

| Protein name | Species | Accession number |
|---|---|---|
| SbPPase | *Sorghum bicolor* | XP_002452886.1 |
| ZmPPase | *Zea mays* | ACG27183.1 |
| OsPPase | *Oryza sativa* Japonica Group | XP_015626515.1 |
| TaPPase | *Triticum aestivum* | XBH80502.1 |
| RcPPase | *Rosa chinensis* | XP_024166692.1 |
| CsPPase | *Cannabis sativa* | XP_030485133.1 |
| VvPPase | *Vitis vinifera* | XP_059591955.1 |
| MoPPase | *Melissa officinalis* | TRINITY_DN13913 |
| ZoPPase | *Zingiber officinale* | XP_042417704.1 |
